# Choice of friction coefficient deeply affects tissue behaviour in epithelial vertex models

**DOI:** 10.1101/2022.11.07.515433

**Authors:** Pilar Guerrero, Ruben Perez-Carrasco

**Affiliations:** Grupo Interdisciplinar de Sistemas Complejos, Departamento de Matemáticas, Universidad Carlos III de Madrid, 28911 Leganés, Madrid, Spain; Department of Life Sciences, Imperial College London, South Kensington Campus, London, United Kingdom

**Keywords:** Vertex model, Cell cycle, Friction coefficient, Tissue mechanics, Epithelial mechanics, Cell cycle variability

## Abstract

To understand the mechanisms that coordinate the formation of biological tissues, the use of numerical implementations is necessary. The complexity of such models involves many assumptions and parameter choices that result in unpredictable consequences, obstructing the comparison with experimental data. Here we focus on vertex models, a family of spatial models used extensively to simulate the dynamics of epithelial tissues. Usually, in the literature, the choice of the friction coefficient is not addressed using quasi-static deformation arguments that generally do not apply to realistic scenarios. In this manuscript, we discuss the role that the choice of friction coefficient has on the relaxation times and consequently in the conditions of cell cycle progression and division. We explore the effects that these changes have on the morphology, growth rate, and topological transitions of the tissue dynamics. These results provide a deeper understanding of the role that an accurate mechanical description plays in the use of vertex models as inference tools.

## 1. Introduction

The spatiotemporal organization of epithelial cells is central to many biological processes. During embryo development, the formation and growth of epithelial layers orchestrate the axes formation and establish the cell lineages in the embryo [1, 2, 3, 4, 5, 6, 7, 8, 9, 10, 11, 12]. In addition, the mechanics of epithelial structures play a major role in tumor growth and malignancy [13]. The geometrical simplicity of epithelia has allowed the development of many mathematical models to understand the tissue’s morphology. Capturing the relationship between the biophysical properties of individual cells and tissue-level properties [14, 15, 16, 17, 10], such an approach provides a powerful tool to acquire generalistic insight into the tissue.

Unfortunately, direct application of analytical frameworks to particular biological tissues is many times not possible. The complexity resulting from the highly dynamic biological processes involved, such as the coupling of cell mechanics and cell cycle progression, makes necessary the development of numerical tools capable of recapitulating the growth of the tissue [7]. In this aspect, computational models have become an indispensable tool to predict and distinguish between hypothesized biophysical mechanisms, as well as to plan future experiments. One particular family of approaches that have proven to be highly successful to model epithelial tissues is that of vertex models [18, 19, 20, 21, 5]. These models assume packing geometries in which cell sheets are approximated by tessellations of polygons [22, 15, 23, 2]. Under this prescription, cells can be fully described as a set of vertices. Tissue dynamics can be naturally incorporated by formulating the evolution of the vertices of the cells as an energy minimization problem [15] resulting in the relaxation of a prescribed Hamiltonian. The flexibility of this approach allows to include of tailored biophysical energetic terms accounting for biophysical properties of the cells such as adhesion and contractibility. Additionally, it also enables the inclusion of cell-specific physiological processes such as cell cycle progression, differentiation, or apoptosis.

Traditional approaches describe the tissue morphology through the quasi-steady state solution of the Hamiltonian [14, 24, 15]. Nevertheless, this queasy-steady solution does not allow to include active deformations of the tissue. Thus, a formulation of the equations of motion dictating the evolution of the tissue is necessary. The most adopted solution to this problem is to assume that all the vertices follow overdamped dynamics. Collectively, cells passively respond to the resulting forces in a viscoelastic manner. Hence, to reveal the mechanics of tissue morphogenesis, we must understand the relationship between active force generation and passive viscoelastic response at the multicellular level. While this bestows the model with the ability to reproduce more realistic dynamics, the effect that different choices of viscosity have on the resulting dynamics is rarely explored [25]. The rationale behind this lies in the quasi-static approaches that assume that the main features of the tissue are determined by equilibrium conditions. In this paper, we study the influence that different choices for the friction coefficient of the overdamped response have on tissue growth, making emphasis on the feedback between mechanical effects and cell cycle duration.

## 2. Vertex dynamics

In order to describe the evolutionary dynamics of the characteristic polygonal morphology of a planar vertex model, the forces induced by the prescribed energy potential can be encoded into an equation of motion for the *N* vertices describing the tissue **r** = {*r*_*i*_|*r*_*i*_ ∈ ℝ^2^, *i* = 1, …, *N*}. The dynamics of every single vertex can be described as a deterministic over-damped equation of motion [24], where inertial terms are neglected compared to dissipative terms, and the friction parameter γ is constant for every single vertex in the system,

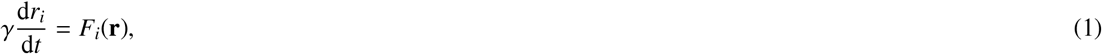

where *F*_*i*_ denotes the total force acting on vertex *i* at time *t*, and it can be directly derived from the energy potential *E* as *F*_*i*_ = − ∇_*i*_*E* (see Fig. 1 A). The choice for the energy potential includes the biophysical properties of the cell relevant for the particular tissue under study and usually has the form [15],

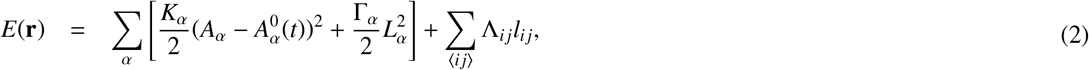

where the first sum runs over all the cells α in the tissue incorporating cell-intrinsic energy terms of elasticity and contractility. The elasticity depends on the deviation from an autonomous preferred area 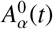 modulated by an elasticity constant *K*_α_. Similarly, the contractility of the cell is introduced as an elastic term that depends on the perimeter of the cell *L*_α_ and a contractility constant Γ_α_. The second sum in Eq. (2) runs over all the edges ⟨*i j*⟩ incorporating cell interaction terms such as cell-cell adhesion energy, where Λ_*i j*_ is a constant representing the line tension. More sophisticated expressions for the potential can include other effects such as inhomogeneities of the contractile tension or non-harmonic energy terms.

**Fig. 1.**
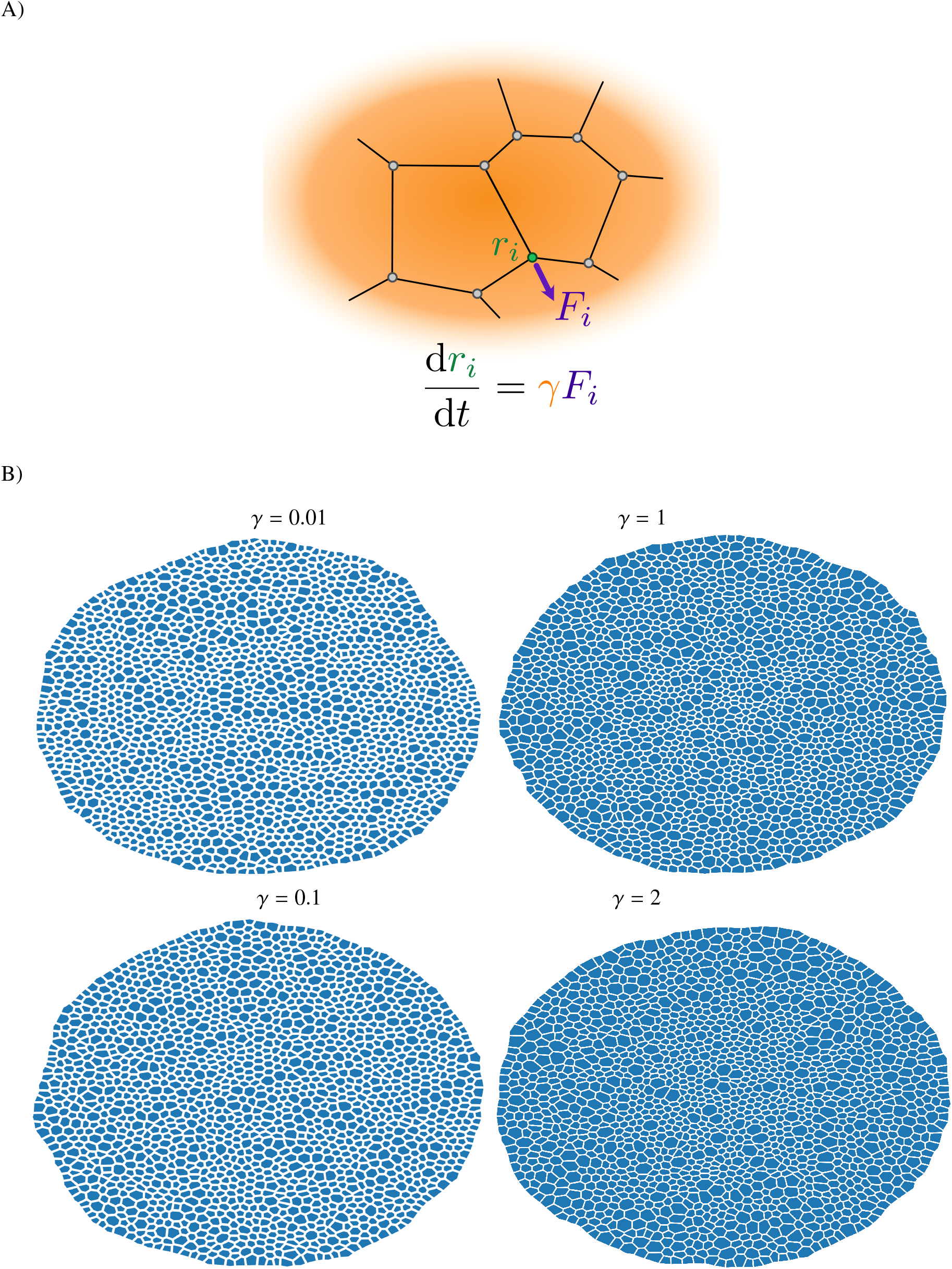
A) Force scheme applied in each vertex of a cell. B) Tissue plot for different values of the friction γ : 0.01 (red), 0.1 (purple), 1 (blue) and 10 (green). Simulation duration *t* = 1600 ≃ 5*T*_*c*_. Other parameter values are provided in Table 1.

## 3. Effect of friction coefficient on cell morphology

The prescribed equations of motion given by Eqs. (1,2) allow a reparametrization of time *τ* = *t*/*γ* that suggests that the role of the friction coefficient *γ* does not perturb the expected configuration of the tissue. Under this assumption, the analysis of the morphology of the tissue is usually reduced to understand the roles of adhesion and contractility [15, 23, 2, 26]. Unfortunately, this dynamical symmetry breaks when other timescales influence the behaviour of the tissue. The main mechanism by which this happens is cell proliferation in which cells divide and change their morphology as they progress through the cell cycle. This is included in vertex models through a time dependence in the energy term (usually encoded by a change in the optimal cell area *A*_0_(*t*)). This active deformation of the cell not only will directly affect its shape but will also affect the topology of the network when cell divisions occur. How relevant are these effects in the epithelium configuration will depend on the relationship between the different timescales.

In addition to cell-autonomous time scales, it is common to assume that there are physical constraints to cell-cycle progression such as the requirement for the cell to reach a certain critical size, *A*_*c*_ and a minimum cell-cycle time of average *t*_*c*_ for cytokinesis to take place [15, 27, 28, 26, 12, 29]. All these factors introduce feedback between the timescales of relaxation and cell cycle progression obscuring the effect that the choice of *γ* has on the epithelium morphology. In order to explore these different effects we made use of the vertex model implementation TifoSi [29]. Specifically, in order to unravel the mechanic’s feedback on the cell cycle progression we used the simplified prescribed target area,

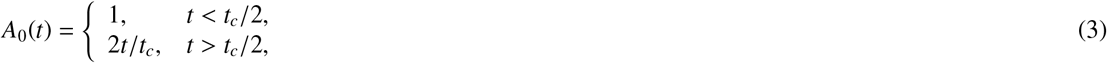

where *t*_*c*_ is the intrinsic cell cycle length of the cell. To incorporate variability in cell cycle progression among different cells, *t*_*c*_ is described as a stochastic variable assigned to each cell at birth. We described *t*_*c*_ as a sum of a deterministic minimum cell cycle duration and a stochastic exponential event [29],

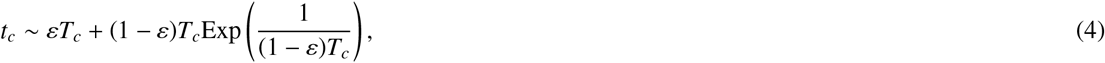

being *T*_*c*_ the average threshold time required for a cell to divide (⟨*t*_*c*_⟩ = *T*_*c*_), and *ε* ∈ [0, 1] a parameter that weights the contribution of the deterministic and the stochastic part. Since *t*_*c*_ is assigned to each cell independently of the interaction with other cells, we will refer to this variability as *intrinsic*. A summary of the values used throughout the manuscript can be found in Table 1, other details can be found in the Supplementary Material.

**Table 1.** Simulation parameter values. If not stated explicitly, the values are the same as the ones in [29].

| Parameter | Description | Value u.a. |
| --- | --- | --- |
| $K$ | Elasticity coefficient | 1 |
| $\Lambda$ | Line tension coefficient | 0.05 |
| $\Gamma$ | Cell perimeter contractility | 0.02 |
| $A_f$ | Maximum target area of a mitotic cell | 2 |
| $A_c$ | Division critical area | $0.78A_f$ |
| $T_c$ | Average intrinsic cell cycle duration | 388 |
| $\varepsilon$ | Stochasticity of intrinsic cell cycle duration | 0.8 |

To examine the potential effects that the choice of friction has on the tissue morphology, we compared the resulting epithelia for different values of the friction constant *γ* that start with an identical hexagonal lattice of 100 cells for a duration of *t* = 3200 ≃ 10*T*_*c*_. Visual inspection of the resulting tissues reveals that cell size variation increases with the friction coefficient, see Fig. 1 B.

Further quantitative analysis of these differences shows that despite area heterogeneities, the tissues are geometrically very similar. The distribution of relative areas as a function of a number of neighbours follows the quadratic property of the cell arrangement described in [30] for smaller cells. Discrepancies only appear for larger cells with more than 7 neighbours, see Fig. 2 A). Similarly, the average shape index of the cells defined as the ratio between the apical perimeter of the cell and the square root of its area remained independent of the friction value chosen, suggesting that the measured change in cell area across friction values is isotropic, see Fig. 2 B). In addition, the shape index has been identified in previous studies with the change of phase in the tissue [31, 32]. For the parameters explored we obtained values below the solid-liquid transition, suggesting that the friction of the tissue did not affect the phase behaviour of the tissue that stayed at every moment in a solid (jammed) phase. Exploration of other parameters such as distribution of polygon shapes, cell area at fixed final tissue size, and cell perimeter distributions depict a similar scenario and can be found as part of the Supplementary Material. All in all, the morphology of the cells seems to be conserved for different values of the friction, with the main difference being an overall slight decrease in the size of the cells for increasing values of friction. Specifically, we observed a reduction of 20% of the average apical area over the three decades of values of the friction explored (Fig. 3 A)).

**Fig. 2.**
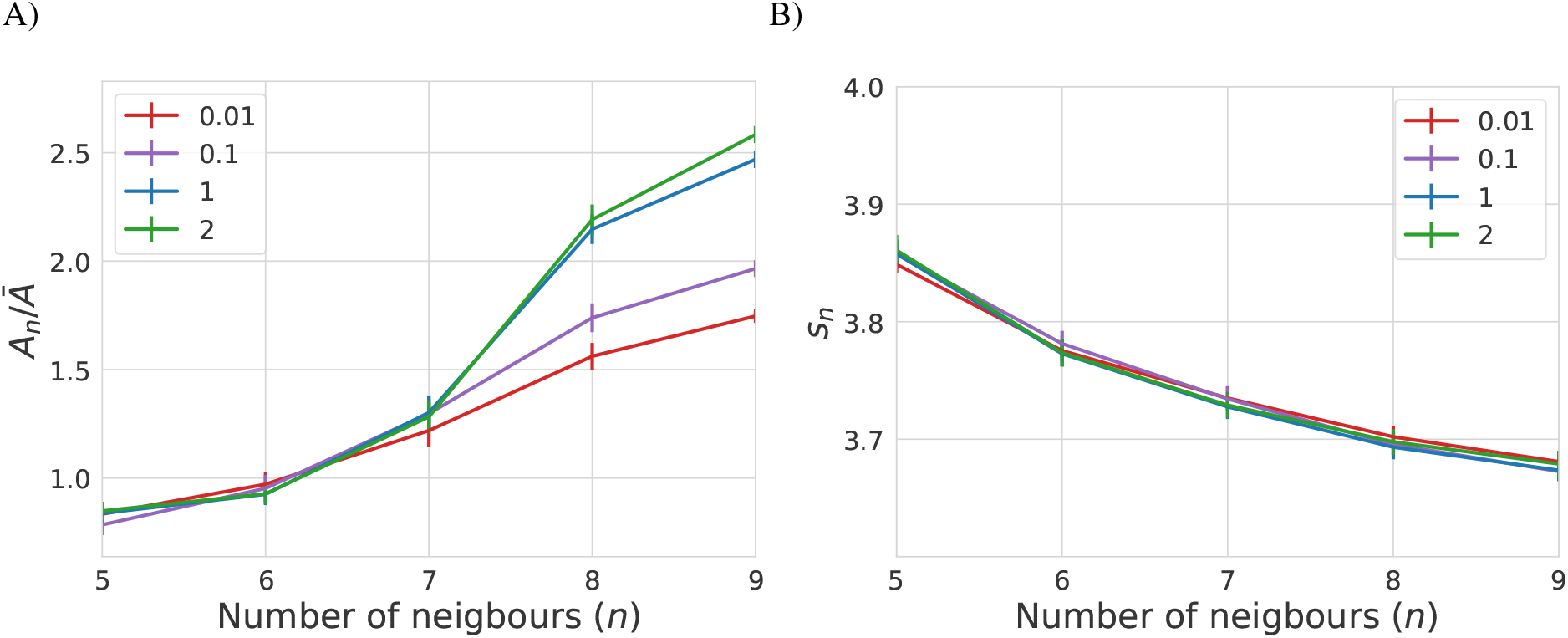
**A) Average cell area *A*_*n*_ normalized by the mean area *Ā* as a function of the number of neighbors, *n*. B) Average shape index, 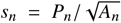, with *P* the perimeter of the cell, as a function the number of neighbours, *n*. Each line is the average of 20 different realizations with different values of *γ* : 0.01 (red), 0.1 (purple), 1 (blue) and 10 (green). Error bars indicate the standard error of the mean (SEM). Simulation duration t = 3200. Other parameter values are provided in Table 1.**

**Fig. 3.**
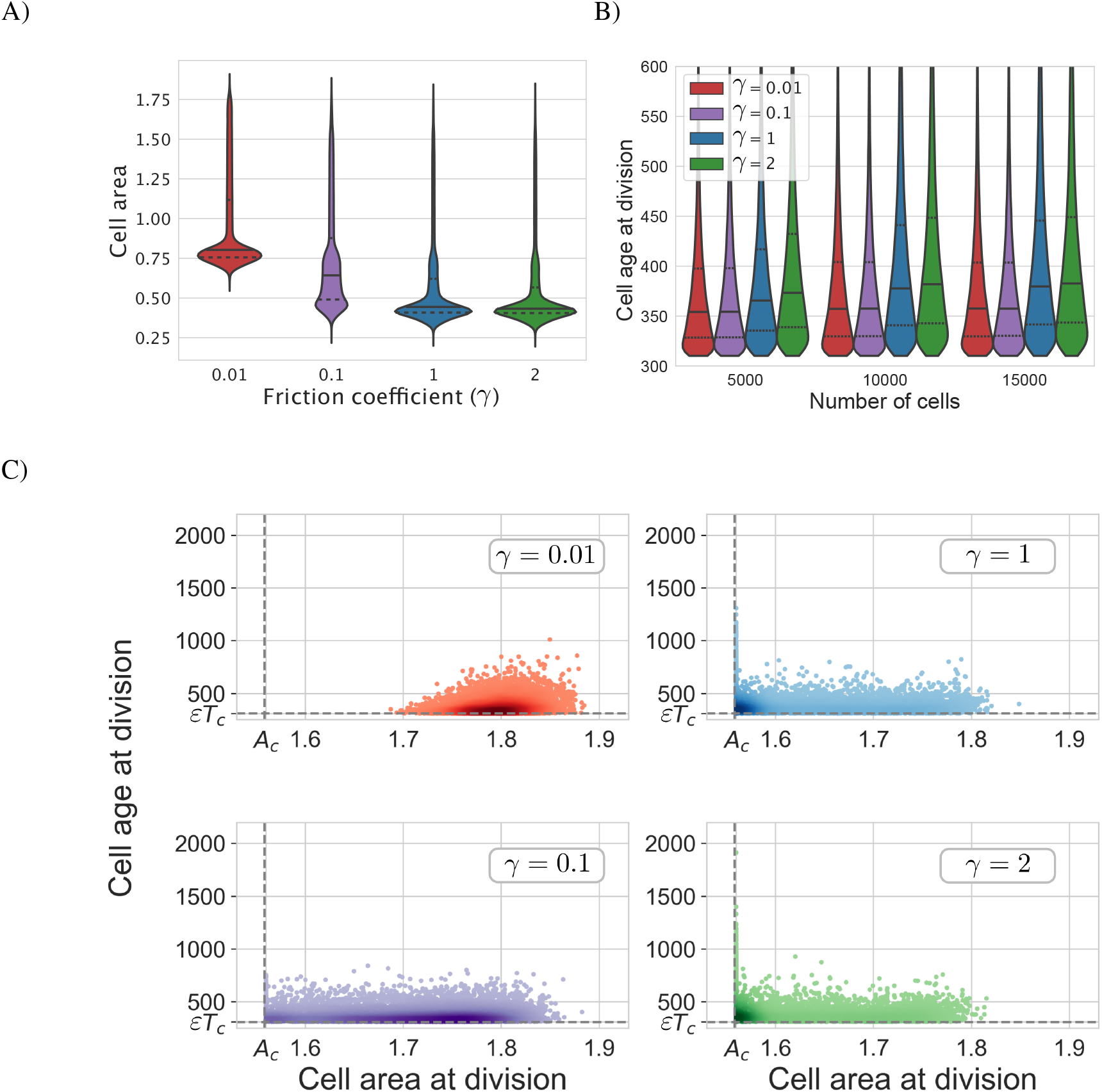
**Variation of cell size and division properties. A) Distribution of cell areas in the tissue aggregated for 20 different simulations at *t* = 3200. B) Division time distribution for different values of *γ* at different tissue sizes. Lines indicate the median (solid line) and quartiles (dotted lines) for the distribution resulting from aggregating 20 different realizations. C) Comparison of cell area and age at division time. *εT*_*c*_ is the minimum duration of any cell cycle (see Eq. 4) and *A*_*c*_ is the critical area required to divide. Simulation corresponds to 20 tissues of 15000 cells aggregated. Parameter values are provided in Table 1.**

## 4. Effect of friction coefficient on tissue growth

The reduction in apical cell area as friction increases can be understood through the relationship between friction and apical cell junction rearrangement required for mitosis. This was confirmed upon inspection of the apical areas of the cells at the moment of division (see Fig. 3 C). Furthermore, analysis of the age of the cell upon division reveals that cells with lower friction values divide faster than high friction ones (see Fig. 3 B). This inverse relationship between size and age of division emerges from the prescribed conditions for cytokinesis. Lower frictions allow for a fast rearrangement of the tissue allowing mitotic cells to reach the minimum required division area *A*_*c*_ before the cell cycle minimum duration *t*_*c*_. On the other hand, for higher frictions, the rearrangement is slower and cells require time beyond *t*_*c*_ to reach the apical cell division area *A*_*c*_. Thus, the tissue friction *γ* controls the interplay between spatial and temporal cytokinesis conditions, attaining the limiting temporal or spatial scenarios for respectively low or high values of *γ*.

This effect that *γ* has on the cell cycle results in a change in the average cell cycle length and consequently a dramatic change in the tissue size at different times (see Fig. 4). In order to check if we can recapitulate the change in tissue growth with a change in the average cell cycle length, we compared the results of the simulations with the deterministic exponential growth that would be expected for the computational average cell cycle duration obtained for each gamma,

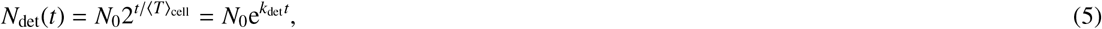

where *N*_0_ is the initial number of cells and ⟨*T*⟩_cell_ is the average cell duration obtained in the simulations. The third expression in Eq. (5) is identical to the second but expressing the growth in terms of the growth rate *k*_det_ ≡ ln(2)/⟨*T*⟩_cell_. Strikingly, the deterministic growth (Eq. (5)) predicts a growth faster than the computational observation (dashed lines in Fig. 4). We hypothesized that not only the change in the mean cell cycle length but also the spread of the distribution of cell cycle durations have a relevant effect on the cell cycle duration and cannot be neglected. For a stationary cell cycle distribution the growth of the tissue will still be exponential but with an effective rate *k*_sto_,

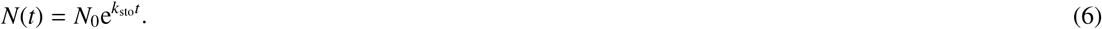

**Fig. 4.**
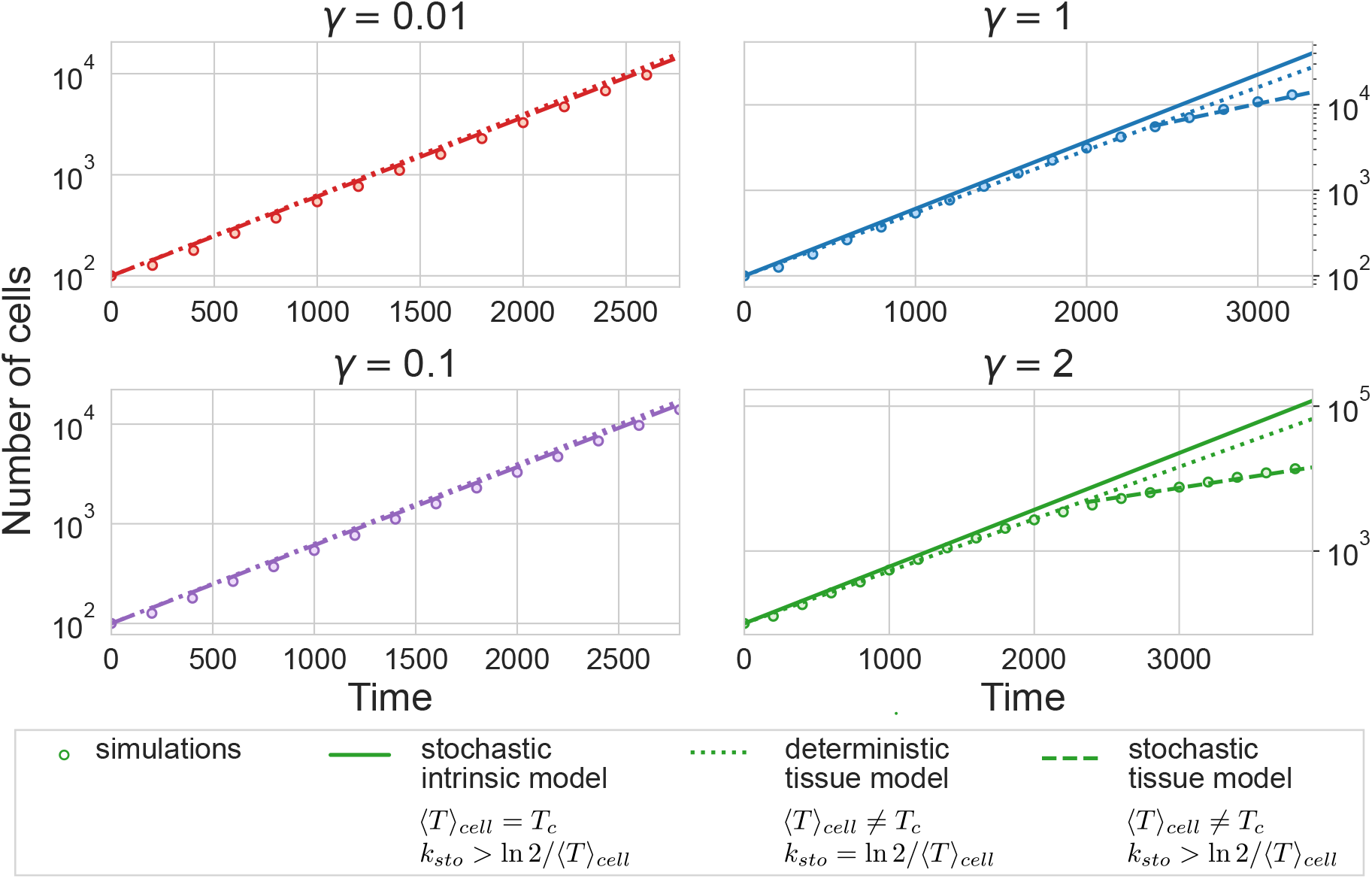
**Comparison of the simulated tissue size (circles) with different deterministic and stochastic predictions (lines) for different values of the friction coefficient. The stochastic intrinsic model (solid line) corresponds to the model without cell-cell interaction, Eq. (15). The deterministic tissue model corresponds with an exponential growth model with duplication time obtained from the average in the tissue (⟨*T*⟩_*cell*_), Eq. (5). The stochastic tissue prediction was obtained by introducing the numerical age structure at the last snapshot of the tissue in Eq. (16). Tissues were simulated until a size of 15000 cells. Other parameter values are provided in Table 1.**

In order to relate *k*_sto_ with the distribution of cell cycle durations we can write a continuity equation for the density of cells 𝒩 at time *t* with an age *τ*, where the age of a cell is the time elapsed since the birth of that cell [33]. The density 𝒩 (*t, τ*) is related to the cell number *N*(*t*) through normalization,

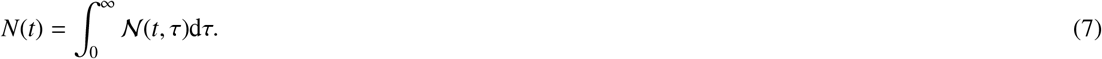

In a small window of time *δ* the density of cells 𝒩(*t, τ*) is reduced an amount proportional to the probability *g*_*δ*_(*τ*) that a cell of age *τ* divides,

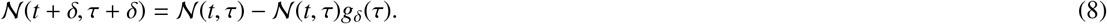

For a given distribution of cell cycle durations *P*(*τ*) that does not change as the tissue grows, similar to the distributions observed in the vertex model simulations, we can write *g*_*δ*_(*τ*) as,

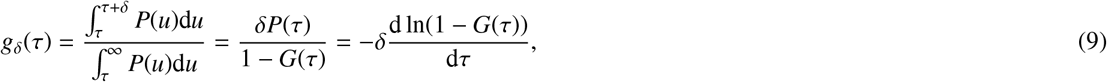

where 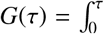 *P*(*t*)d*t* is the cumulative density function of cell cycle durations. Introducing the expression for the division probability (Eq. (9)) in the continuity equation (8) and taking the limit *δ* ⟶ 0 we get the differential form of the continuity equation,

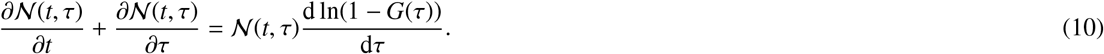

The first term of Eq.(10) can be related to the growth rate of the tissue 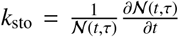. This gives rise to an ordinary differential equation for the age *τ*,

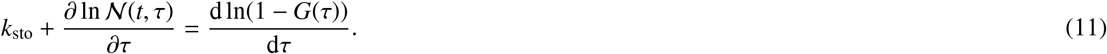

Integrating Eq. (11) we obtain an explicit expression for the age contribution to the cell density 𝒩(*t, τ*),

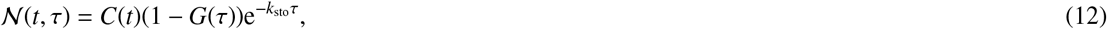

where *C*(*t*) is the integration constant depending on time *t* that corresponds with the density of newly born cells 𝒩 (*t*, 0) = *C*(*t*). This term can be determined by the proliferation boundary condition i.e. every cell finishing the cell cycle gives raise to two newly born cells,

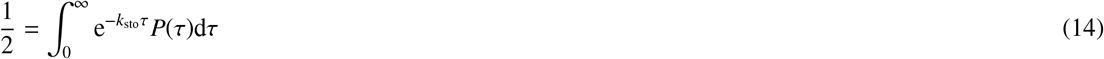

Introducing the expression for the density (Eq. (12)) in the boundary condition (Eq. (13)), we obtain a closed Euler-Lotka equation that relates the rate *k*_sto_ with the cell cycle length distribution,

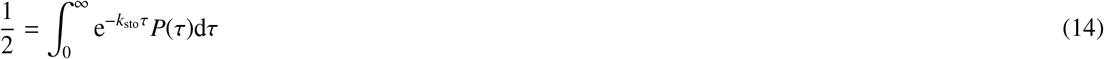

Note that there is not an immediate relationship between *k*_sto_ and *k*_det_, the latter depending exclusively on the first moment of the cell cycle distribution 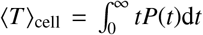. For the intrinsic cell cycle distribution (Eq. (4)), in the absence of cell-cell interaction (⟨*T*⟩)_*cell*_ = *T*_*c*_), Eq. (14) becomes,

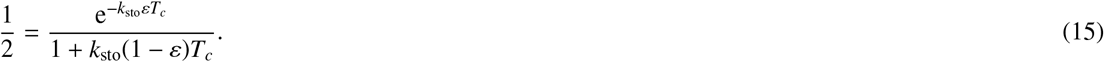

In the deterministic cell cycle scenario (*ε* = 1) we recover the deterministic growth *k*_sto_ = ln 2/*T*_*c*_. On the other hand, for the purely exponential case (*ε* = 0) we obtain an upper bound for the tissue growth rate *k*_sto_ = 1/*T*_*c*_. For intermediate values of *ε*, the rate *k*_sto_ can be obtained by solving numerically the transcendental equation (15). For the parameters used in this paper (see Table. 1) we have *k*_sto_*T*_*c*_ ≃ 0.70, which predicts a tissue growth very close to the deterministic scenario. In general, for the stochastic cell cycle model used Eq. (4) it is expected that the intrinsic stochasticity will increase the value of *k*_sto_ i.e. while preserving the same average cell cycle duration, the tissue will grow at a faster speed than in a purely deterministic scenario, which is the opposite effect to the slow down predicted by the simulation of the vertex model (solid line in Fig. 4).

This apparent mismatch is not surprising, the distribution of cell cycle durations in the simulation will be different from the intrinsic stochastic cell cycle in the absence of cell-cell interactions (cf. Eq. 4 and Fig. 2). Since we cannot access analytically the expression for cell cycle durations in the tissue *P*(*τ*), the actual value *k*_*sto*_ can be calculated by solving numerically the integral equation 14. Alternatively, one can integrate the continuity equation (10) for all the possible ages *τ*, and use the normalization of 𝒩 and the proliferation boundary condition Eq. (13) obtaining the relationship,

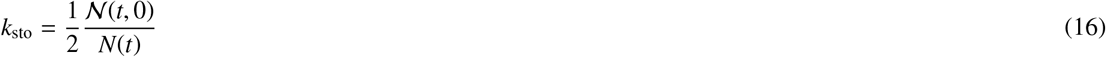

that relates directly the rate of growth of the tissue and the age structure of the population. Comparison of the rates of growth of the tissue with the computational observation is able to recover the discrepancies that appeared with the analysis (dashed line in Fig. 4). This reveals the relevance that cytokinesis conditions impose on the growth of the tissue by shaping the cell cycle length distribution.

### 4.1. Topological transitions are also affected by the friction constant

In the previous section, we showed that friction has a dramatic effect on the growth of the tissue by affecting the prescribed typical mitosis conditions used in the vertex model. In particular, the threshold condition in critical cell area *A*_*c*_ seems to have a relevant impact on the mean duration of the cell cycle and its variability. To explore further this effect we explored the resulting tissues from a model that does not require a threshold in the target area in order to divide (*A*_*c*_ = 0). As expected, the resulting tissue grows faster, following a growth more similar to the intrinsic stochastic growth model Eq. (15) (see Fig. 5 A). Nevertheless, the agreement with the deterministic model is not perfect, suggesting that more factors affect the timely mitosis of the cells.

**Fig. 5.**
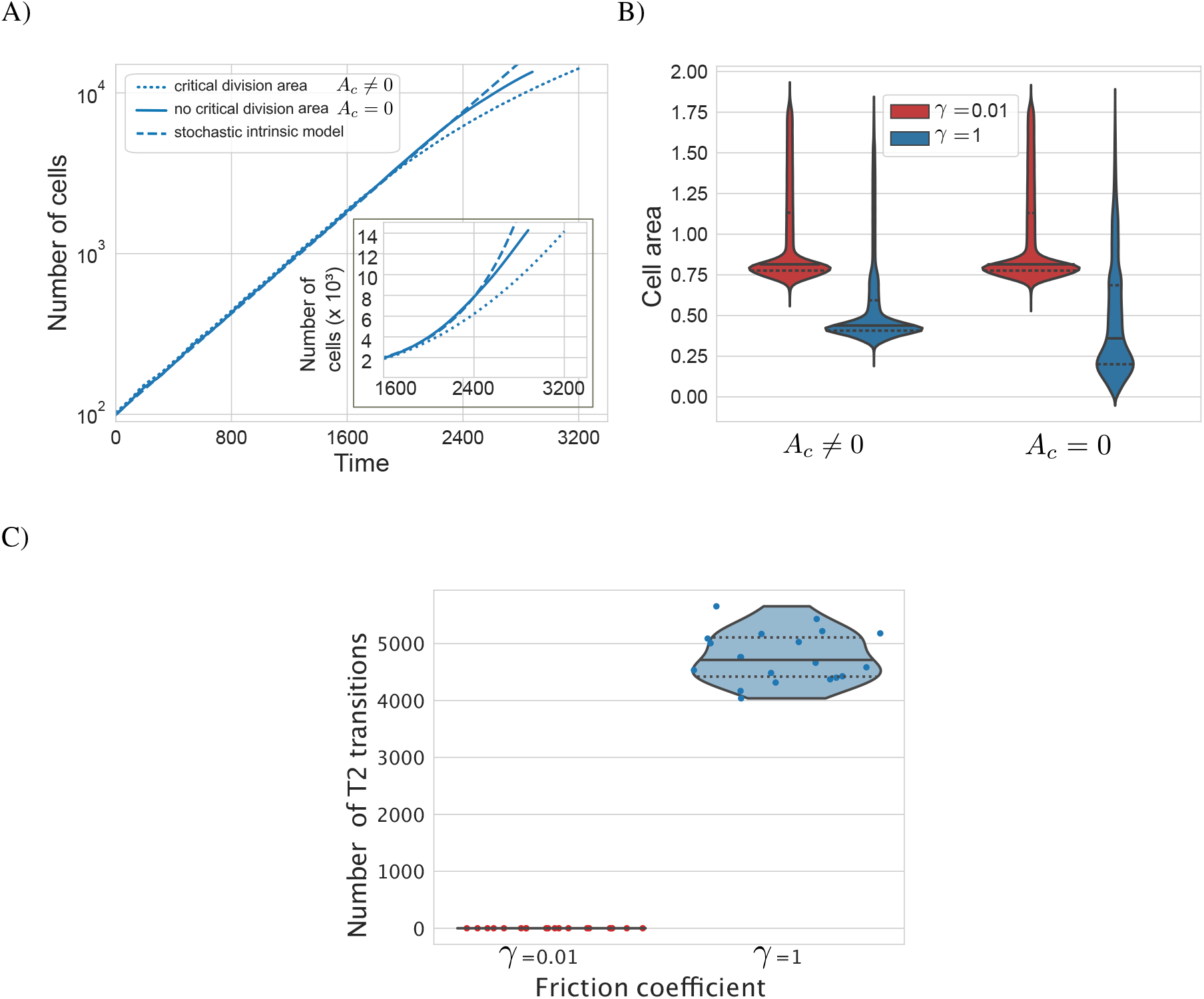
**Effect of topological transitions in tissue growth. A) Comparison of tissue growth with and without critical area conditions for mitosis (dotted and solid lines) for *γ* = 1. Both conditions result in a tissue smaller than expected from the intrinsic growth (dashed line) resulting from Eq. (15). Inset: zoom on the diverging parts of the curves using a linear scale. Results are the average of the 20 simulations for each condition. B) Distribution of cell areas in the tissue at 15000 cells for different values of *γ* and condition of critical division area. Violin plots show the aggregated data from 20 simulations for each condition. C) Number of T2 transition events in the absence of critical area condition for two different values of *γ*. Each conditions is the aggregate of 20 realizations (circles). All violin plots indicate the median (solid line) and the quartiles (dotted lines). Other parameter values are provided in Table 1.**

Analysis of the configuration of cell areas confirms that removing the condition of critical area affects the distribution of cell sizes for large values of *γ* but does not rescue the distribution of cell sizes observed at lower values of *γ* (see Fig. 5 B). Cells do not require any more to reach a certain size to divide, which results in overall smaller cell sizes at division, and consequently smaller cell areas at the start of the cell cycle of both daughter cells. Since in the absence of a cell critical area, cells progress along the cell cycle without affecting each other, the cell area should not affect the timing of division and tissue growth. The only mechanism left by which cells will change their cell cycle progression is a topological transition. Specifically, T2 transitions control the removal of cells when they become too small. This mechanism has been employed in the past to explain apoptosis or extrusion of cells from the epithelium [12]. Analysis of the amount of T2 transitions confirms that the removal of *A*_*c*_ results in an increased number of T2 transitions sufficient to explain the discrepancies between simulations with different friction coefficients (Fig. 5). This is only observed for high values of *γ* where the slow vertex dynamics impede the growth of the cells up to the target area *A*_0_.

## 5. Conclusion

Vertex models are a very powerful computational tool to simulate epithelial organization. One of the advantages is the possibility of incorporating intrinsic mechanistic details of cell cycle progression through a target area term in the Hamiltonian [12, 29]. This allows us to understand the non-linear effects by which cell-intrinsic mechanistic details affect overall tissue dynamics. At the same time, it reveals how sensitive are the results to different modelling choices that are usually taken arbitrarily and regularly in the literature.

To make this evident, in this paper we have focused on the effects of the choice of friction coefficient *γ* in the evolution of the tissue (Eq 1). In particular, we have observed that not only do different frictions result in non-intuitive changes in size and variability of cell areas, but most importantly in tissue growth. Typical prescriptions of the cell cycle in vertex models are limited to incorporating average cell observations in the form of a time-dependent target area and a set of conditions of division. By contrast here we show that the choice of friction will affect dramatically the resulting cell cycle by controlling these division conditions and modifying its mean duration and dispersion. All in all, this reveals the complexity of incorporating intrinsic cell-autonomous mechanistic descriptions into sophisticated vertex models and how accurate descriptions of the variability of cell cycle duration in time are paramount to achieving a useful description of the tissue.

As part of the integration of these non-linear effects in tissue growth, we showed that stochastic distributions of the cell cycle can be tackled through continuity equations for the age of the cells in the tissue. This approach can accommodate also the effect of T2 transitions by incorporating cell clearance along its cell cycle. For instance, an asymmetric division in which only a fraction of daughter cells stays in the tissue after division [34] could be incorporated replacing the factor 2 in Eq. (13). All in all, these calculations highlight the relevance of measuring full distributions of cell cycle lengths in experimental assays as opposed to reporting summary statistics.

The current study has tackled the usual dissipative prescription by which the displacement per unit of time is proportional to the force homogeneously in the tissue (Eq. (1)) where *γ* is a global constant of the tissue. Future studies should also question this approximation that is based on the implicit physical assumption that each vertex is a point of mass with identical assigned friction. By contrast, one can expect that the dissipation of each vertex is related to the total material that needs to be displaced under a certain force e.g. is not the same to drag the membranes joining three small cells, than the membranes of three bigger ones.

All in all, the ultimate goal of vertex models is to be used together with experimental data to understand the mechanistic properties of tissue dynamics. The observations of this paper, together with recent studies revealing a lack of parameter identifiability of vertex models [35], make evident that more work is required in the field in order to exploit reliably vertex models as tools to infer mechanistic properties from experimental data.

## Acknowledgments

P.G. acknowledges Ministerio de Ciencia, Innovación y Universidades/European Regional Development Fund (Spain/EU) grant PGC2018-098186-B-I00 (BASIC). R.P.-C. acknowledges the department of Life Sciences at Imperial College London

