## Supplementary material for "Choice of friction coefficient deeply affects tissue behaviour in epithelial vertex models": SM.pdf

---

---

### Simulation files

All the simulations were produced using the vertex model implementation TiFoSi. An example of the TiFoSi input .xml file used to run the simulations is attached as part of the Supplementary Material. The input .xml file contains all the parameters required to run the simulations with the most important parameters summarized in Table 1 and Eqs. (3)-(4).

### Supplementary Figures

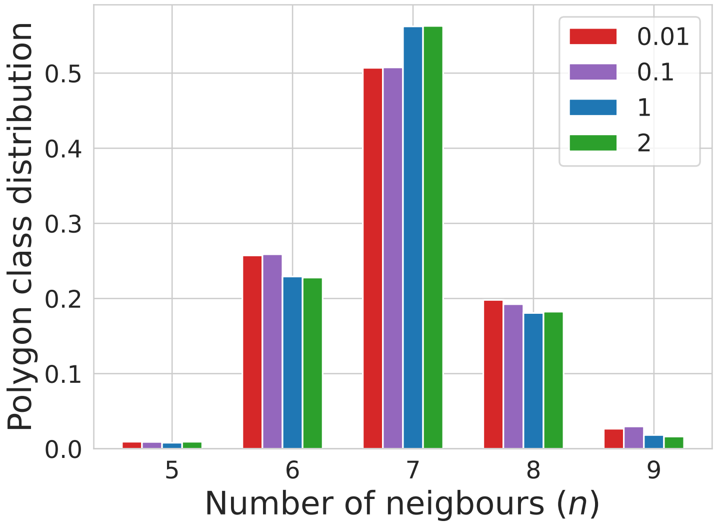


Figure SM.1: Polygon cell distribution for different value for different values of $\gamma$ : 0.01 (red), 0.1 (purple), 1 (blue) and 10 (green). Data aggregated from 20 simulations ended at 15000 cells. Other parameter values are provided in Table 1.

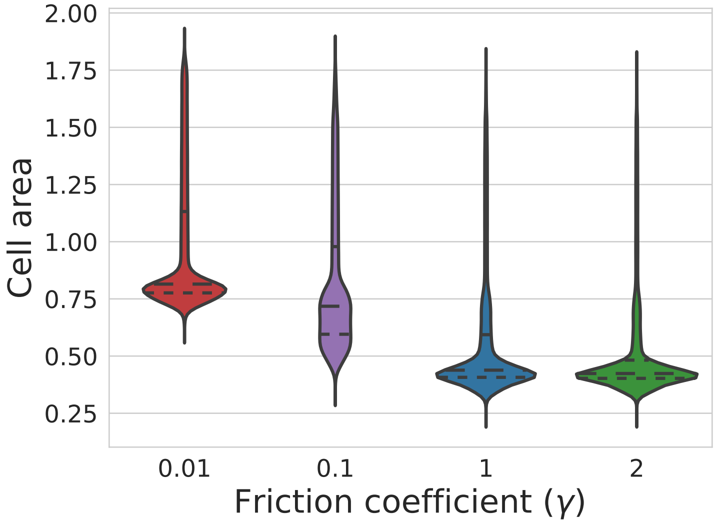


Figure SM.2: Cell area distribution for different values of $\gamma$ : 0.01 (red), 0.1 (purple), 1 (blue) and 10 (green). Data aggregated from 20 simulations ended at 15000 cells. Other parameter values are provided in Table 1.

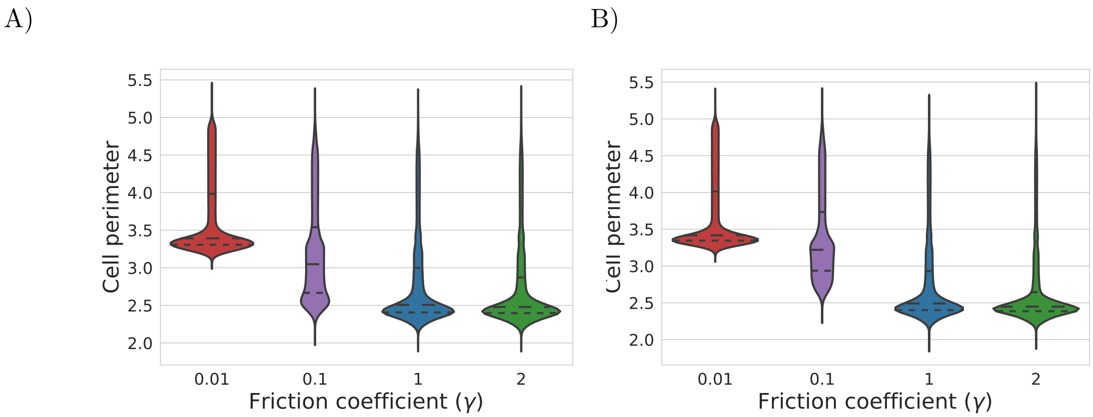


Figure SM.3: Violin plot o cell perimeter distributions for different values of $\gamma$ : 0.01 (red), 0.1 (purple), 1 (blue) and 10 (green). A) Simulation total duration t=3200. B) Simulation ended at 15000 cells. Data aggregated from 20 simulations ended at 15000 cells. Other parameter values are provided in Table 1.
